# Divergence-based introgression polarization

**DOI:** 10.1101/539197

**Authors:** Evan S. Forsythe, Daniel B. Sloan, Mark A. Beilstein

**Affiliations:** Department of Biology, Colorado State University, Fort Collins, CO 80523, USA.; School of Plant Sciences, University of Arizona, Tucson, AZ 85721, USA.

## Abstract

Introgressive hybridization results in the transfer of genetic material between species, often with fitness implications for the recipient species. The development of statistical methods for detecting the signatures of historical introgression (IG) in whole-genome data has been a major area of focus. While existing techniques are able to identify the taxa that exchanged genes during IG using a four-taxon system, most methods do not explicitly distinguish which taxon served as donor and which as recipient during IG (i.e. polarization of IG directionality). The existing methods that do polarize IG are only able to do so when there is a fifth taxon available and that taxon is sister to one of the taxa involved in IG. Here, we present *Divergence-based Introgression Polarization* (*DIP*), a method for polarizing IG using patterns of sequence divergence across whole genomes, which operates in a four-taxon context. Thus, *DIP* can be applied to infer the directionality of IG when additional taxa are not available. We use simulations to show that *DIP* can polarize IG and identify potential sources of bias in the assignment of directionality, and we apply *DIP* to a well-described hominin IG event.

## INTRODUCTION

Hybridization is an influential evolutionary force (Stebbins 1968) that is widespread in natural populations (Yakimowski and Rieseberg 2014; Mallet et al. 2016). Through backcrossing to parental populations (Rieseberg and Soltis 1991), hybrids can serve as bridges for the transfer of alleles and adaptive traits between species or populations (Rieseberg and Soltis 1991; Dasmahapatra et al. 2012; Suarez-Gonzalez et al. 2016), a process known as introgression (IG) (Rieseberg and Soltis 1991; Rieseberg et al. 1996; Green et al. 2010; Dasmahapatra et al. 2012; Mallet et al. 2016). Whole genome sequences and advances in phylogenetic methods (Soltis and Soltis 2003) have revealed signatures of historical IG in scientifically and economically important groups, including well-studied examples in Neanderthals and non-African human populations (Kuhlwilm et al. 2001; Green et al. 2010). Several methods have been developed to identify taxa that exchanged genes during IG (Huson et al. 2005; Than et al. 2008; Green et al. 2010; Durand et al. 2011; Martin et al. 2015; Pease and Hahn 2015; Stenz et al. 2015; Rosenzweig et al. 2016). While these methods generally perform well across a variety of biological and experimental scenarios (Zheng and Janke 2018), theoretical and empirical work have identified conditions under which each method is susceptible to bias (Eriksson and Manica 2012; Rosenzweig et al. 2016).

While there are many tools for detecting IG between taxa, a more challenging aspect of IG analyses is identifying taxa serving as donors *vs.* recipients of genetic material during IG (i.e. IG directionality). If hybrids successfully backcross to both parents during IG, alleles will move in both directions, meaning each parent will serve as donor for some introgressed loci and recipient for other loci. However, if backcrosses with one parent but not the other are favored by physiological (Rieseberg and Soltis 1991), selective (Orive and Barton 2002), or biogeographical (Currat et al. 2008) factors, it can lead to asymmetrical (Barton and Hewitt 1985) movement of alleles (directional IG, denoted hereafter with ‘⇒’). IG has been shown to underlie the transfer of adaptive traits to recipient lineages (Whitney et al. 2006; Dasmahapatra et al. 2012; Dannemann et al. 2016; Figueiró et al. 2017), so the ability to infer the directionality of IG (i.e. polarize IG) is essential in order to form hypotheses about the functional and adaptive consequences of IG.

The majority of tests to detect the occurrence of IG do not explicitly polarize IG (Zheng and Janke 2018), and those that can only do so in certain cases. For example, the *D* statistic (Green et al. 2010) is widely-used to infer instances of IG in a four-taxon system. IG polarization is possible under *D* only when data for a fifth taxon are available (Green et al. 2010; Pease and Hahn 2015). Moreover, the fifth taxon must be sister to one taxon involved in IG but cannot itself be involved in IG. Pease and Hahn (2015) define this specific configuration of introgressing taxa and sister taxa as “intergroup” IG and describe how, when these specific five-taxon conditions are met, the branching order of introgressed gene trees indicates directionality. However, the authors also describe how other types of IG (e.g. “ancestral” IG) cannot be polarized. There are many cases in which a fifth taxon with the required phylogenetic placement is either not sampled or does not exist (Forsythe et al. In Review). In these cases, it is possible to statistically identify IG using existing methods but not necessarily to polarize IG. Thus, there is a need for a more widely applicable statistical method to distinguish between bidirectional and unidirectional IG, while identifying donor and recipient taxa.

Here, we describe and test a method for inferring directionality of IG from genome-scale data, which we refer to as *Divergence-based Introgression Polarization* (*DIP*). *DIP* is based on the observation that, when IG occurs, it alters not only the level of nucleotide sequence divergence between the two species exchanging genes (Rosenzweig et al. 2016) but also divergences with related species that are not directly involved in IG; these changes occur in systematic and predictable ways according to the directionality of IG (Fig. 1) (Forsythe et al. In Review; Fontaine et al. 2015; Hibbins and Hahn, In Review). *DIP* is calculated from pairwise sequence divergence between taxa involved in IG and a sister taxon, comparing divergence values obtained from introgressed loci *vs*. non-introgressed loci. It takes as input the same types of data used to infer IG by existing methods (whole genome/chromosome alignments or single-gene alignments of loci sampled throughout the genome). However, unlike existing methods, *DIP* can polarize IG when only four taxa are sampled, meaning *DIP* is more widely applicable than existing methods.

**Fig. 1.**
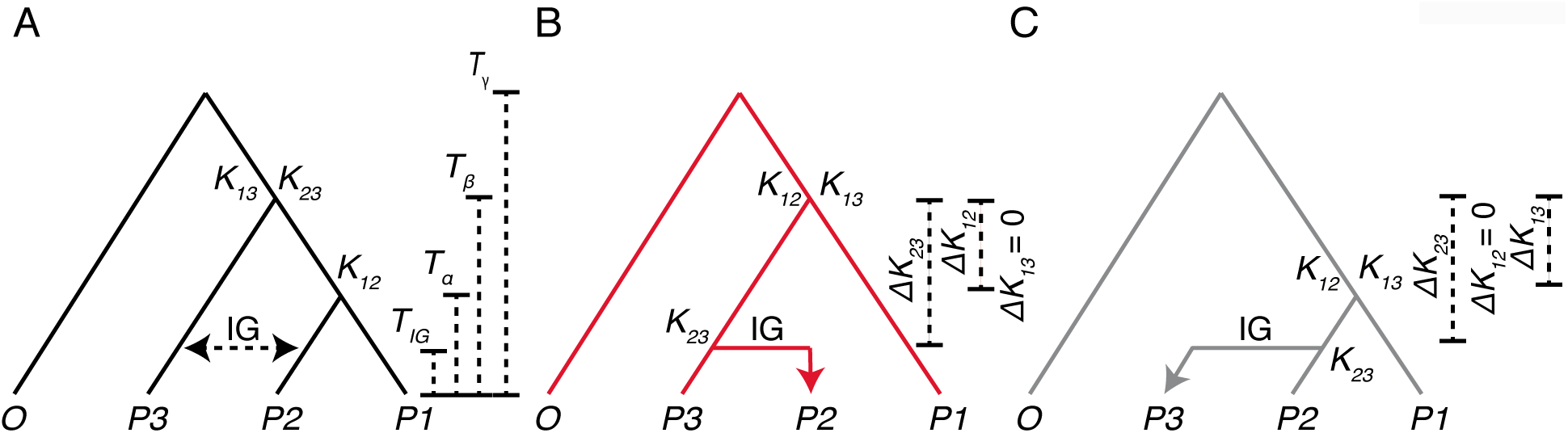
Expected divergence under simulated introgression. The species *P1, P2, P3*, and *O* were used for simulation analyses. (**A**) The species branching order (SP). IG between species *P2* and *P3* is indicated with a double-sided dotted arrow. Default values used during all simulations, unless specified otherwise, are: *T*_*IG*_=1, *T*_*α*_*=*4, *T*_*β*_=8, and *T*_*γ*_=12 in coalescent units (4*N* generations) (Hudson 2002). (**B**) A gene tree depicting a gene that was introgressed *P3*⇒*P2*. (**C**) A gene tree depicting a gene that was introgressed *P2*⇒*P3*. *ΔK* values are calculated based on changes in mean divergence between pairs of taxa in the set of SP trees *vs.* the set of IG trees (see *Eq. 1-3*). Note that the expected profiles of *ΔK* values for *P3*⇒*P2* IG differs from that of *P2*⇒*P3* IG, forming the basis for the *DIP* test (see Main Text and Fig. 2).

We present tools to implement the *DIP* method: https://github.com/EvanForsythe/DIP. We also simulate whole genome alignments in which a subset of loci were introgressed either unidirectionally, asymmetrically, or symmetrically. We use these simulated genome alignments to assess how accurately *DIP* polarizes asymmetrical IG and to investigate the effects of parameters that are known to affect existing IG inference methods, such as proportion of IG and timing of IG (Durand et al. 2011; Martin et al. 2015; Zheng and Janke 2018). We have recently used the principles of *DIP* to document asymmetrical IG among Brassicaceae species (Forsythe et al. In Review), and here, we also apply *DIP* to an empirical data from modern and archaic hominins.

## NEW APPROACHES

IG can alter levels of sequence divergence between taxa, and these changes can differ depending on the directionality of IG (Forsythe et al. In Review; Hibbins and Hahn, In Review) (Fig. 1). To define the properties of a divergence-based IG test, we use hypothetical species *P1, P2, P3* and an outgroup, *O*. Species *P1* and *P2* are sister within the species tree, and we model IG between species *P2* and *P3*. We denote the timing of the three successive speciation events among these taxa as *T*_*γ*_, *T*_*β*_, and *T*_*α*_ and the timing of the IG event between *P2* and *P3* as *T*_*IG*_ (Fig. 1A). When introgression has occurred between *P2* and *P3,* some loci will reflect a history of IG, while other loci will reflect a history of speciation. In applying *DIP*, a gene tree is inferred for each locus, and the resulting topology is used to distinguish introgressed loci (IG loci) from speciation loci (SP loci). For all loci, we quantify pairwise sequence divergence values between *P2* and *P3* (*K*_*23*_), between *P1* and *P2* (*K*_*12*_), and between *P1* and *P3* (*K*_*13*_) (Fig. 1). The values of *K*_*23*_, *K*_*12*_, and *K*_*13*_ on a given gene tree are expected to correspond to *T*_*IG*_, *T*_*α*_, and *T*_*β*_ in a way that depends on the IG history of that gene. IG in either direction is expected to reduce *K*_*23*_ relative to genes that reflect the species tree, as the divergence time between the sequences of these taxa is reduced from *T*_*β*_ to *T*_*IG*_ (Fig. 1). In contrast, IG can cause *K*_*12*_ to increase corresponding to a change in divergence time from *T*_*α*_ to *T*_*β*_ but only if IG occurred from *P3* to *P2* (Fig. 1B). IG in the other direction should not affect *K*_*12*_. The effects on *K*_*13*_ are also sensitive to the direction of IG. If it occurs from *P2* to *P3*, IG should decrease *K*_*13*_ based on a change in divergence time from *T*_*β*_ to *T*_*α*_ (Fig. 1C), but there should be no effect on *K*_*13*_ if IG occurs in the other direction. To quantify these effects, differences are calculated between the mean values of *K*_*23*_, *K*_*12*_, and *K*_*13*_ from all SP loci and the mean values of the same corresponding divergence measurements from all IG loci in the following fashion:

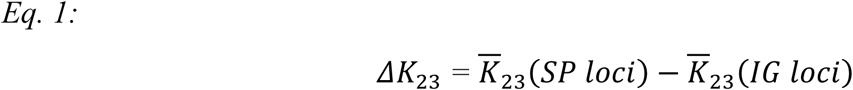

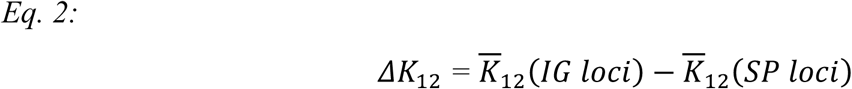

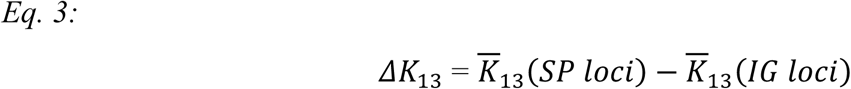

Note that the order of subtraction used in defining these terms is not always the same with respect to SP and IG loci and was chosen such that the effects of relevant IG are expected to yield positive (rather than negative) *ΔK* in each case. Together, this set of *ΔK* values composes the divergence profile of *DIP*. Below we show the relative magnitudes of these values can be used differentiate evolutionary histories based on the polarity of IG. We also use coalescent-based simulations to identify biases that can be introduced by other sources of genealogical discordance such as incomplete lineage sorting (ILS), and we devise additional layers of *DIP* comparisons that can be used to partially alleviate these biases.

## RESULTS

### DIP: Distinguishing modes of unidirectional and bidirectional introgression

The simplest application of *DIP* is related to the approach we recently applied in analyzing IG among Brassicaceae species (Forsythe et al. In Review). It involves testing whether *ΔK*_*23*_, *ΔK*_*12*_, and *ΔK*_*13*_ are significantly greater than zero and compares these results to the expectations for *ΔK* under different IG scenarios (Fig. 2). If IG has occurred in both directions between *P2* and *P3*, then all three *ΔK* values should be positive. However, as noted above, if IG has occurred exclusively in one direction, the expectation for either *ΔK*_*12*_ or *ΔK*_*13*_ should remain zero (Fig. 2). To test the performance of *DIP*, we simulated whole-genome alignments under unidirectional IG in each direction, as well as under symmetric bidirectional IG (see Methods and Fig. S1). We applied *DIP* to each simulated genome. For the genome simulated under unidirectional *P2⇒P3* IG, we observed *ΔK*_*23*_ > 0, *ΔK*_*12*_ = 0, and *ΔK*_*13*_ > 0 (Fig. 3A), which is the expected pattern for that direction of IG (Fig. 1). For the genome simulated under symmetric bidirectional IG, we observed *ΔK*_*23*_ > 0, *ΔK*_*12*_ > 0, and *ΔK*_*13*_ > 0 (Fig. 3B), which is the expected pattern if some IG is occurring in both directions. For the genome simulated under unidirectional *P3⇒P2* IG, we observed *ΔK*_*23*_ > 0, *ΔK*_*12*_ > 0, and *ΔK*_*13*_ = 0 (Fig. 3C), again reflecting our expected *DIP* profile for that direction. These results indicate that *DIP* can correctly classify all three types of IG under these simulated conditions.

**Fig. 2.**
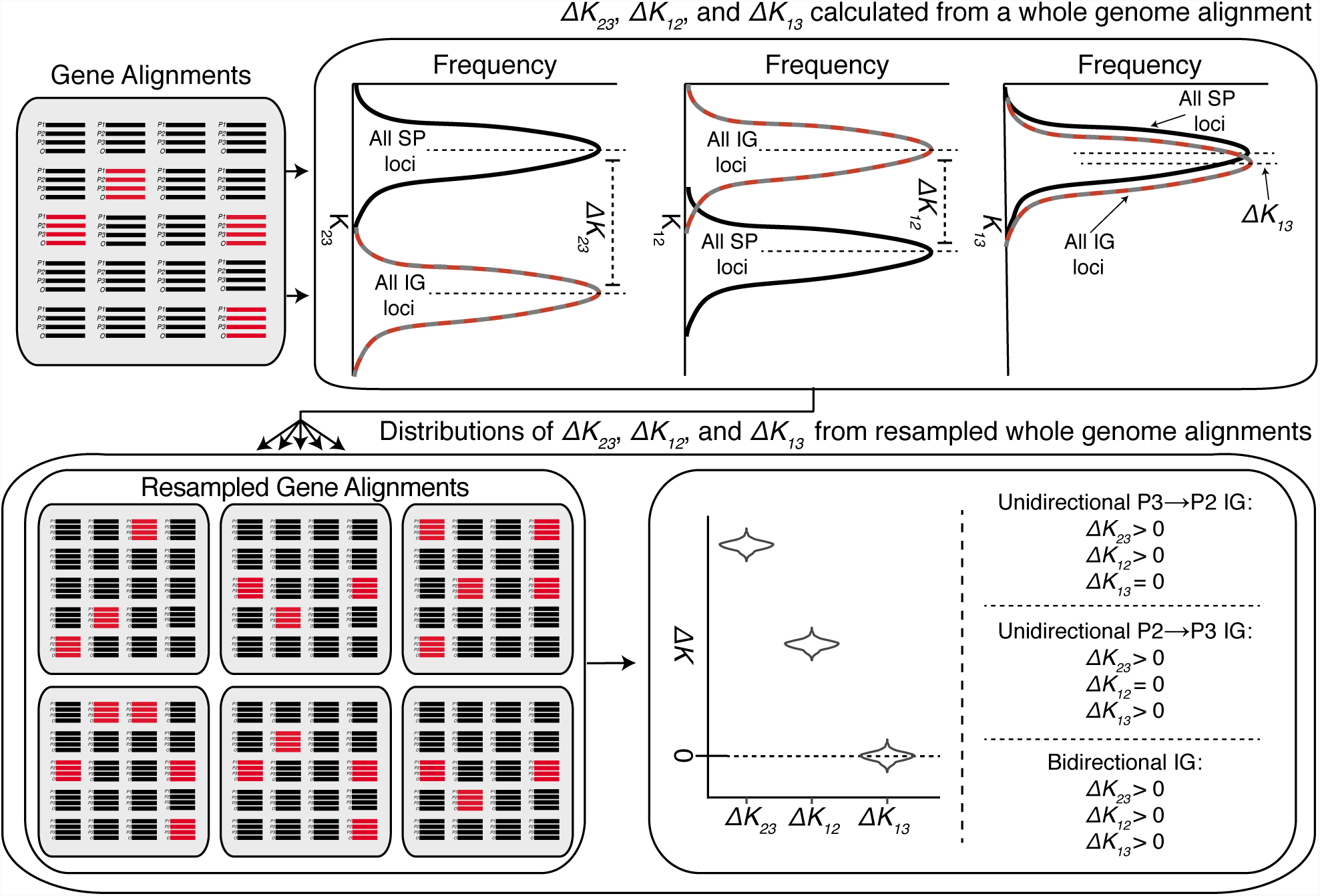
Workflow of the *DIP* test. Point estimates of *ΔK*_*23*_, *ΔK*_*12*_, *ΔK*_*13*_ are calculated from whole genomes, which are then resampled to yield distributions of *ΔK*_*23*_, *ΔK*_*12*_, *ΔK*_*13.*_ Unidirectional *P3⇒P2* IG is indicated by the profile, *ΔK*_*23*_ > 0, *ΔK*_*12*_ > 0, and *ΔK*_*13*_ = 0. Unidirectional *P2⇒P3* IG is indicated by *ΔK*_*23*_ > 0, *ΔK*_*12*_ = 0, and *ΔK*_*13*_ > 0. Bidirectional IG is indicated by *ΔK*_*23*_ > 0, *ΔK*_*12*_ > 0, and *ΔK*_*13*_ > 0. All other profiles are considered inconclusive regarding the occurrence and directionality of IG. *P*-values for testing whether each *ΔK* value significantly differs from 0 are obtained from the proportion of replicates for which *ΔK* ≤ 0. Colors reflect the black, red, and gray genealogical histories from Fig. 1. In this illustration, all IG loci are in the *P3⇒P2* (red) direction. But we use the red/gray dashed lines for showing the distribution of IG loci because, in general, the set of IG loci can contain *P3⇒P2* loci, *P2⇒P3* loci, or both.

**Fig. 3.**
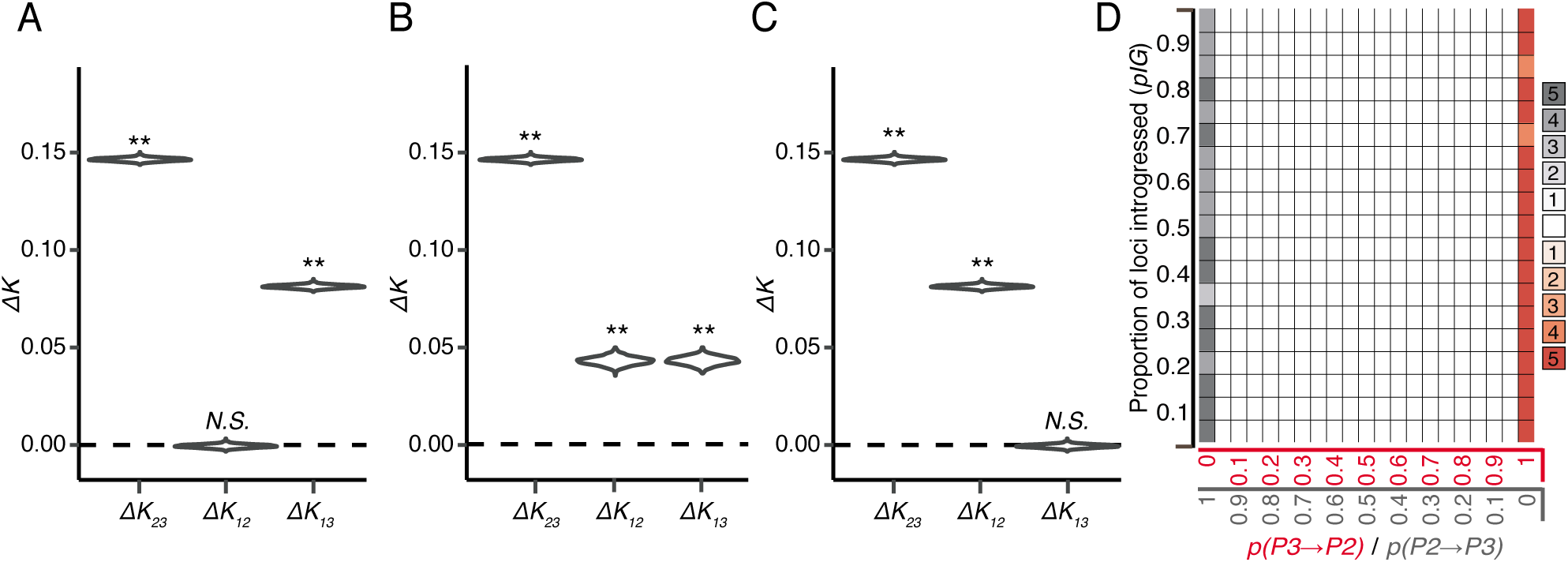
*DIP* analysis of simulated introgression. Genomes were simulated according to steps 1-3 in Fig. S1, under unidirectional *P2⇒P3* IG (**A**), symmetrical bidirectional *P3*⇔*P2* IG (**B**), and unidirectional *P3⇒P2* IG (**C)**. Simulation parameters are as follows: (**A**), *n* = 5000, *pIG* = 0.5, *p(P3⇒P2*) = 0; (**B**), *n* = 5000, *pIG* = 0.5, *p(P3⇒P2*) = 0.5; (**C**), *n* = 5000, *pIG* = 0.5, *p(P3⇒P2*) = 1. *DIP* was applied to each genome to yield profiles of *ΔK*_*23*_, *ΔK*_*12*_, *ΔK*_*13*_. ** indicates significant departure from 0 (*p* < 0.01). (**D**) A plot scanning simulation parameters, proportion of the genome that was introgressed (*pIG*) (y-axes) and proportion of introgressed loci transferred in each direction (*p(P3⇒P2*)) (x-axis). Each square in the plot indicates the *DIP* results obtained from five replicated simulated genome alignments. Red boxes indicate the profile consistent with *P3*⇒*P2* IG (see panel C). Gray boxes indicate the profile consistent with *P2*⇒*P3* IG (see panel A). The shading of the boxes corresponds the number of replicates (out of five) that indicate a given profile, as specified by the key to the right of the plot. Unshaded boxes indicate all five replicates yield the bidirectional IG profile (see panel B).

Next, we explored the performance of *DIP* across a range of different parameter settings, including the proportions of genes in the genome that had been subject to IG (*pIG*). We also varied the proportions of IG loci that moved in one direction or the other [*p(P3⇒P2*)]. We performed a parameter scan (Fig. S1) by generating simulated genomes with different values of *pIG* and *p(P3⇒P2*) and applying *DIP* to each genome (Fig. 3D). We found the expected *P3⇒P2 DIP* profile for the majority of replicated genomes generated with *p(P3⇒P2*)=1 (i.e. unidirectional *P3⇒P2* IG) (Fig. 3D, red boxes). Further, we found the expected *P2⇒P3 DIP* profile for the majority of replicated genomes generated with *p(P3⇒P2*)=0 (i.e. unidirectional *P2⇒P3* IG) (Fig. 3D, gray boxes). Intermediate *p(P3⇒P2*) values all yielded the expected *DIP* profile for bidirectional IG for all replicates (Fig. 3D, white boxes).

### Double-DIP: Detecting asymmetry in cases of bidirectional introgression

Existing IG polarization methods tend to assume unidirectionality of IG, but it is also important to consider the possibility of asymmetric bidirectional IG that falls short of being strictly unidirectional [discussed in (Martin et al. 2015)]. The basic implementation of *DIP* described above can detect the presence of bidirectional IG (see Fig. 3B profile and Fig. 3D white boxes), but it does not report directional asymmetry (i.e. whether either of the two directions predominates) at intermediate values of *p(P3⇒P2*). Hereafter, we refer to this basic implementation of *DIP* as Single-*DIP* or *1*×*DIP*. To more directly test for asymmetry in cases of bidirectional IG, we developed an additional step in the *DIP* analysis, which we refer to as Double-*DIP* or *2*×*DIP*. The premise of *2*×*DIP* is that *ΔK*_*12*_ for loci introgressed *P3*⇒*P2* and *ΔK*_*13*_ for loci introgressed *P2*⇒*P3* have the same expected values, as they are both based on a shift in divergence time between *T*_*β*_ and *T*_*α*_ (Fig. 1). Therefore, under symmetric bidirectional (*P3*⇔*P2*) IG, we expect genome-wide values of *ΔK*_*12*_ and *ΔK*_*13*_ to equal each other. Alternatively, if *P3*⇒*P2* IG exceeds *P2*⇒*P3* IG, we expect genome-wide *ΔK*_*12*_ > *ΔK*_*13*_. *2*×*DIP* compares the magnitudes of *ΔK*_*12*_ and *ΔK*_*13*_ by formulating a simple summary statistic, *ΔΔK*, which is defined as follows:

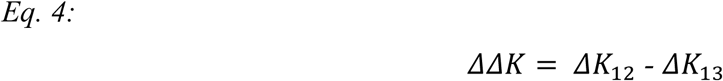

The expectation for the *ΔΔK* summary statistic is zero under symmetric bidirectional IG, positive under IG that is biased towards *P2*, and negative under IG that is biased towards *P3* (Fig. 4).

**Fig. 4.**
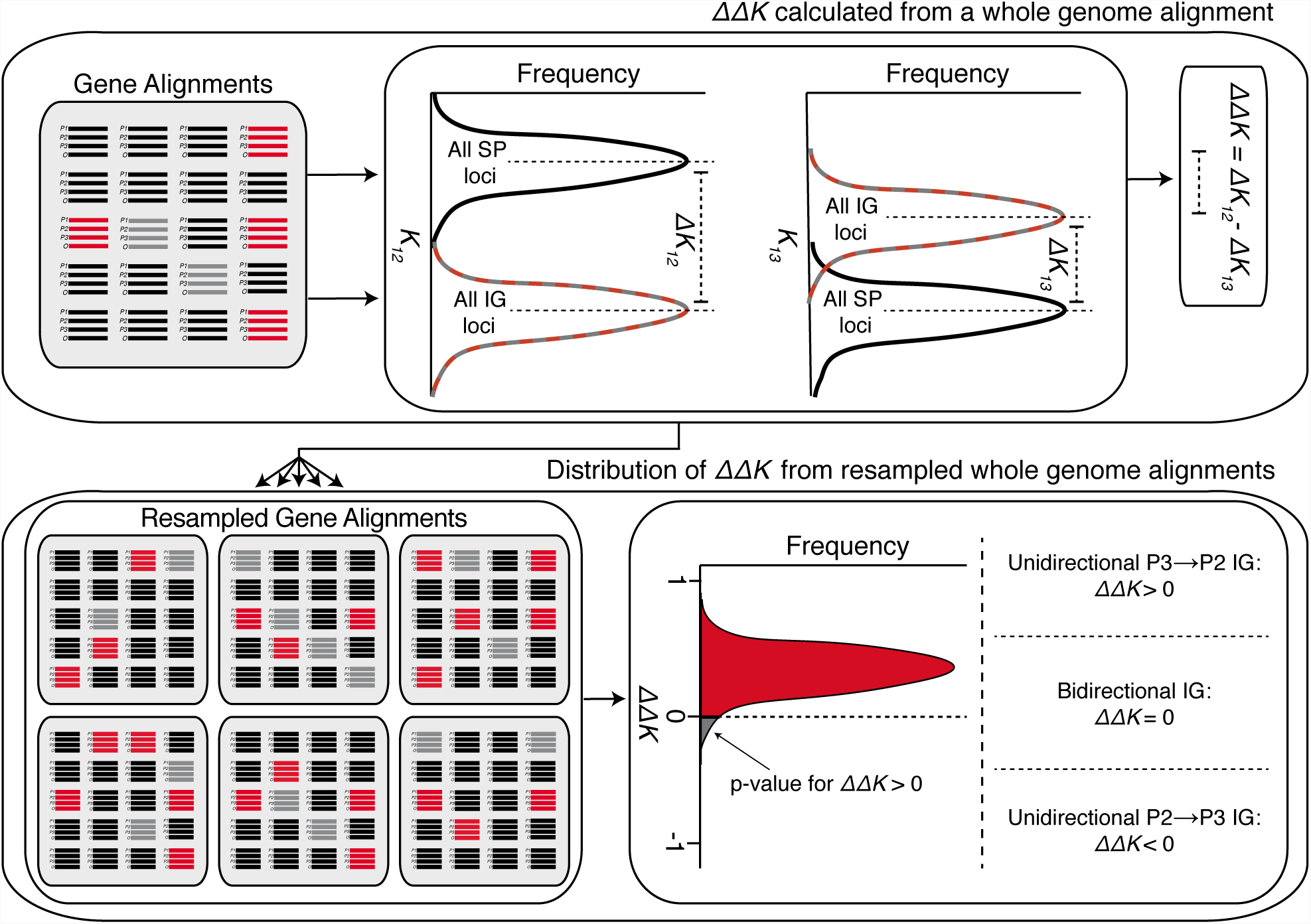
Workflow of the *2*×*DIP* test. (**Top**) A point estimate of *ΔΔK* is calculated from a whole genome alignment from *ΔK*_*12*_ and *ΔK*_*13*_ values. (**Bottom**) A sampling distribution of *ΔΔK* is calculated from resampled gene alignments (bootstrapping) obtained from the original genome. If the majority of *ΔΔK* replicates are > 0, it is an indication of asymmetric *P3⇒P2* IG. In this case, the proportion of *ΔΔK* replicates < 0 determines the *p*-value (doubled for a two-sided test) for asymmetric *P3⇒P2* IG. Asymmetric *P2⇒P3* IG is indicated by the opposite pattern.

We explored the performance of *2*×*DIP* by simulating genomes in the same manner as described above for *1*×*DIP*. For the genome simulated under unidirectional *P2⇒P3* IG (*p(P3⇒P2*) = 0), we observed a significantly negative *ΔΔK* (Fig. 5A, *p <* 0.0002), consistent with our expectations. For the genome simulated under symmetric bidirectional IG, *ΔΔK* did not significantly differ from zero (Fig. 5B, *p =* 0.914), also consistent with expectations. For the genome simulated under unidirectional *P3⇒P2* IG (*p(P3⇒P2*) = 1), we observed significantly positive *ΔΔK* (Fig. 5C, *p* < 0.0002), again reflecting expectations. These results indicate that *2*×*DIP* correctly classified all three types of IG. As above, we also performed a parameter scan to explore *2*×*DIP*. We found that genomes simulated with *p(P3⇒P2*) = 0.5 (i.e. symmetric bidirectional IG) returned *ΔΔK* value that did not significantly differ from zero (Fig. 5D, white boxes). We also found significant *ΔΔK <* 0 for nearly all replicated genomes simulated with *p(P3⇒P2*) < 0.5 and significant *ΔΔK >* 0 for nearly all replicated genomes simulated with *p(P3⇒P2*) > 0.5 (Fig. 5D). The only exception to these patterns were found at *pIG* ≤ 0.1 during nearly symmetrical IG (*p(P3⇒P2*) = 0.45 and 0.55). Taken together, these results indicate that *2* × *DIP* correctly inferred asymmetrical IG, even in cases in which there is only slight asymmetry, meaning it is a sensitive method for polarizing asymmetrical IG that is robust across a wide variety of parameter values.

**Fig. 5.**
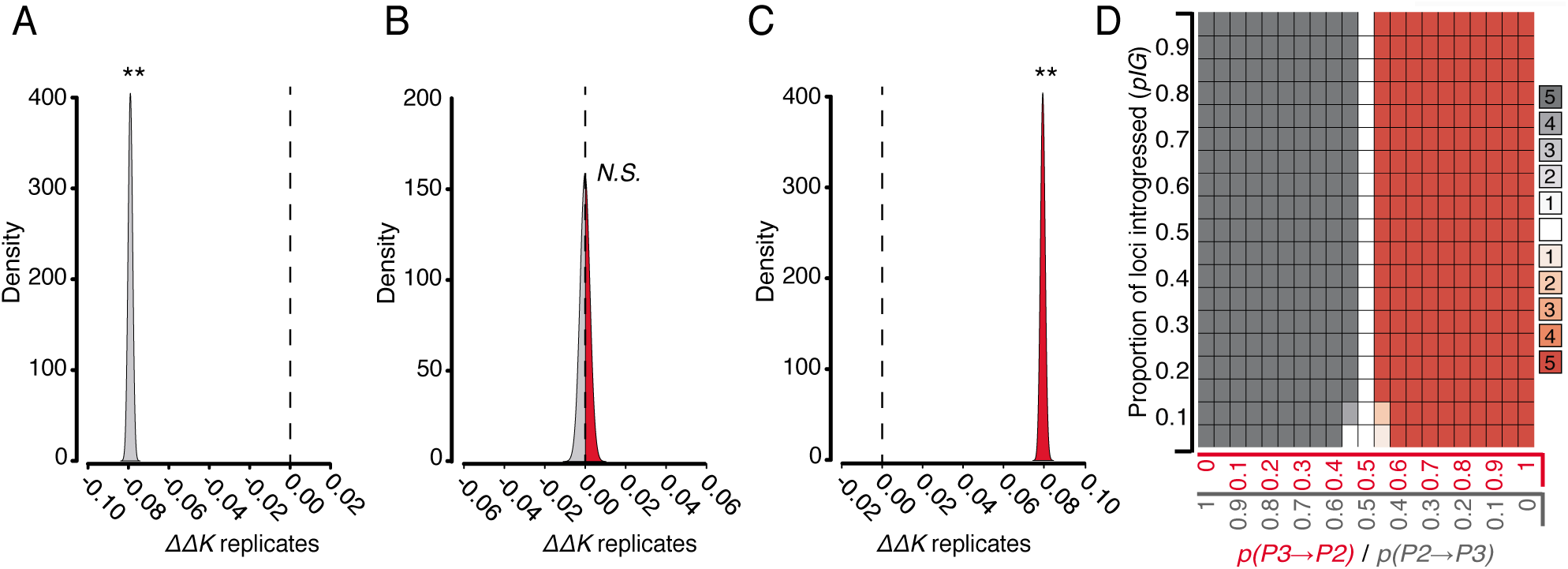
*2*×*DIP* analysis of simulated introgression. Genomes were simulated according to steps 1-3 in Fig. S1. Genomes were simulated under unidirectional *P2⇒P3* IG (**A**), symmetrical bidirectional *P3*⇔*P2* IG (**B**), and unidirectional *P3⇒P2* IG (**C)**. Simulation parameters are as follows: (**A**), *n* = 5000, *pIG* = 0.5, *p(P3⇒P2*) = 0; (**B**), *n* = 5000, *pIG* = 0.5, *p(P3⇒P2*) = 0.5; (**C**), *n* = 5000, *pIG* = 0.5, *p(P3⇒P2*) = 1. *2*×*DIP* was applied to each genome to yield a sampling distribution of *ΔΔK.* ** indicates significant departure from 0 (*p* < 0.01). (**D**) A plot scanning *pIG* and *p(P3⇒P2*) as in Fig. 3D. Red boxes indicate significant (*p*<0.05) *P3*⇒*P2 2*×*DIP* signature (see panel C). Gray boxes indicate significant (*p*<0.05) *P2*⇒*P3 2*×*DIP* signature (see panel A). The shading of the boxes corresponds the number of replicates (out of five) that significantly indicate the signature, as specified by the key to the right of the plot. Unshaded boxes indicate all five replicates failed to reject symmetrical IG (see panel B).

### Robustness of DIP to population divergence time

The task of assigning gene trees as IG *vs.* SP based on gene tree topology is an integral part of *DIP;* however, this task comes with challenges. Phylogenetic methods rely on diagnostic synapomorphies to infer gene tree topologies; scarcity of synapomorphies in an alignment leads to phylogenetic error and inaccurate gene tree assignment. Another confounding factor is ILS, which can result in gene trees that reconstruct the history of deep coalescence, as opposed to the underlying history of SP/IG. ILS can result in introgressed loci displaying the SP topology and vice versa. Importantly, ILS is also expected to yield gene trees displaying an alternative third topology (Green et al. 2010) (see Triple*-DIP* below). Both mis-assignment and ILS are more pronounced during rapid divergence (i.e. short internal branches) and can be investigated with coalescent simulations (Degnan and Rosenberg 2009; Degnan and Rosenberg 2013). Moreover, it has been shown that, because *P3⇒P2* IG trees have longer internal branch lengths than *P2⇒P3* IG trees, the latter are more prone to both mis-assignment and ILS (Zheng and Janke 2018). This feature introduces the potential for directional bias in *DIP*. Therefore, we explored divergence times, as an additional parameter that may influence performance.

All previous simulations were implemented with constant and large divergence times (see Fig. 1). To explore the branch length parameter, we modified divergence times by multiplying all of the branch lengths by a scaling factor (SF) (see Methods), essentially modifying the height of the entire tree used for simulations. SFs greater than one yield taller trees, while SFs less than one yield shorter trees. For each SF, we simulated ten replicate genomes and calculated *ΔΔK* for each replicate. We first classified SP and IG loci based on the known history used to simulate the data and plotted the resulting *ΔΔK* values (omniscient *2*×*DIP*). We found that *2*×*DIP* correctly inferred asymmetry (or lack thereof) at all branch lengths and that the magnitude of *ΔΔK* was proportional to the SF (Fig. 6A, D and G). However, when working with real datasets it is rare to know if individual loci with IG topologies are the result of bona fide IG, as opposed to ILS or errors in phylogenetic inference. To explore the impact of the SF on the ability of *2*×*DIP* to distinguish between signature from bona fide IG loci and those mis-assigned due to ILS or phylogenetic error, we calculated *ΔΔK* using topology-based (non-omniscient) assignment. With this approach, we observed an upward bias in *ΔΔK* at low SFs (Fig. 6B, E, and H). This bias favors inference of *P3⇒P2* IG even when there is asymmetry in the opposite direction (Fig. 6E). As expected, this bias exists at the SFs for which mis-assignment of gene trees is most pronounced (Fig. S2), suggesting that it results from gene tree mis-assignment and/or ILS (see Discussion).

**Fig. 6.**
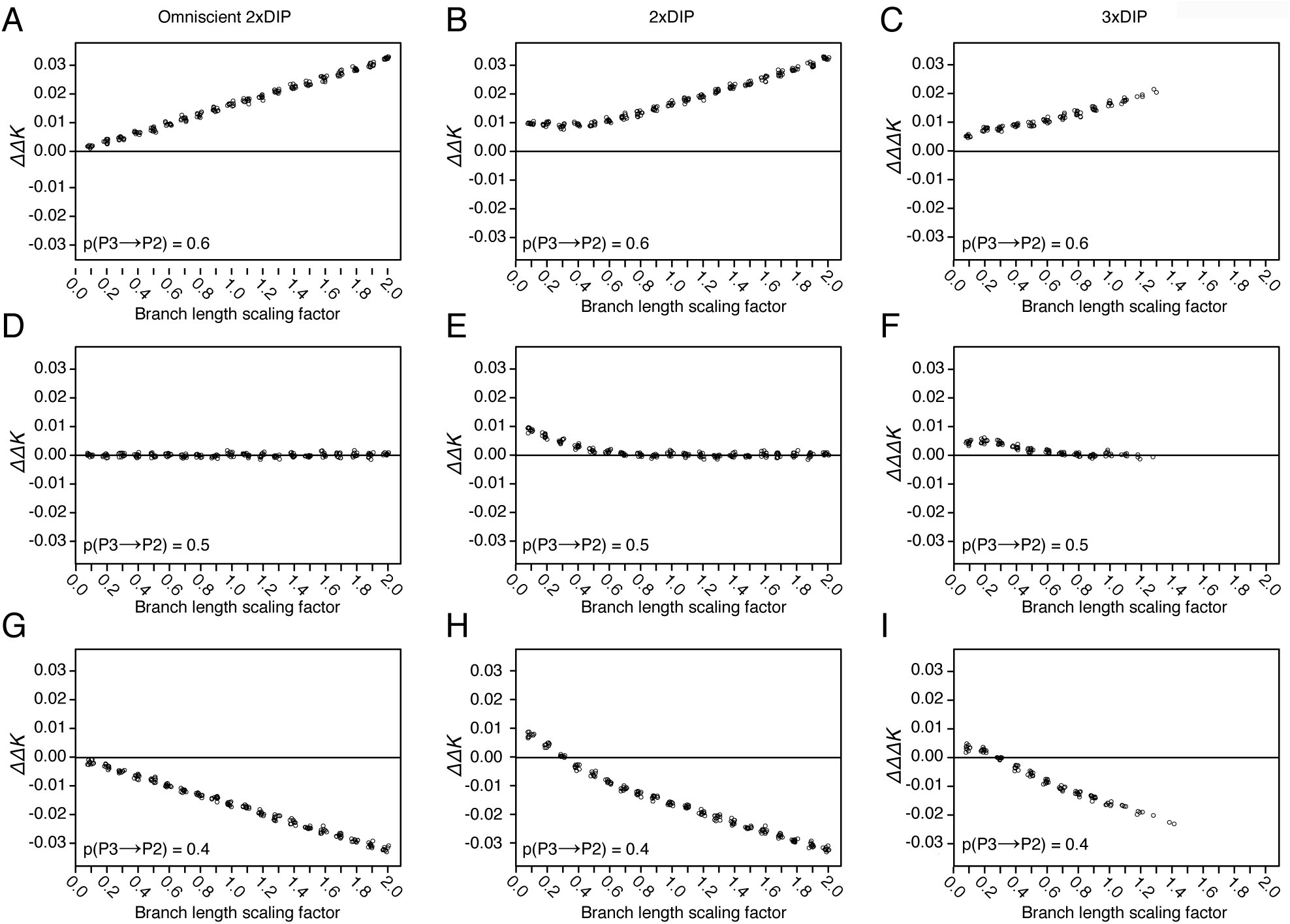
Exploration of branch length parameters used during genome simulation. The default branch lengths used during all previous simulations (*T*_*IG*_=1, *T*_*α*_*=*4, *T*_*β*_=8, and *T*_*γ*_=12) were multiplied by branch-length scaling factors. For all plots, 10 replicate genomes were simulated for each scaling factor value. *pIG* = 0.5 was used for all simulations. *DIP* was performed on each replicate; individual points on plots represent point estimates of *ΔΔK* and *ΔΔΔK* (jittered for clarity). Genomes were simulated with asymmetric IG favoring *P3⇒P2* (**A-C**), symmetric bidirectional IG (**D-F**), and asymmetric IG favoring *P2⇒P3* (**G-I**). Omniscient *2*×*DIP* (**A, D, and G**), standard *2*×*DIP* (**B, E, and H**), and *3*×*DIP* (**C, F, and I**) were performed. *ΔΔΔK* data points are absent at higher scaling factors because this adjusted version of *ΔΔK* can only be calculated when there are at least some loci with the unexpected topology (ALT loci) as a result of topology mis-assignment or ILS.

### Triple-DIP: Adjusting for gene tree assignment bias

To address the directional bias in *2*×*DIP* caused by gene tree mis-assignment/ILS at short branch lengths, we developed an additional layer that can be applied in *DIP* analysis, which we refer to as Triple-*DIP* or *3*×*DIP*, so named because it includes an additional *Δ* component (i.e. the “delta of the delta of the delta”). Briefly, in addition to calculating the standard *2*×*DIP* as above, we also calculate an alternative *ΔΔK* (*ΔΔK*_*alt*_) that substitutes gene trees with the alternative topology, ((*P1, P3*), *P2*), for the IG loci used in the standard *ΔΔK:*

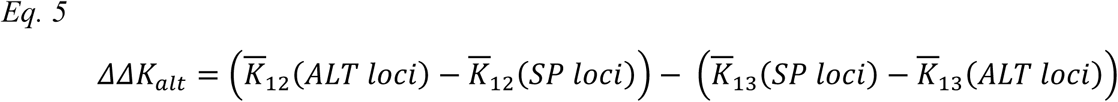

Because *P2* and *P3* are the two taxa subject to IG, loci with this alternative topology should arise only from mis-assignment/ILS and not true IG. Following the logic of standard *D* statistics (Green et al. 2010; Durand et al. 2011), we reasoned that mis-assignment/ILS should be equally likely to produce each of the two topologies that conflict with the species tree. Therefore, this alternative *2*×*DIP* calculation may provide a measure of the amount of bias that is introduced by these processes. In applying *3*×*DIP,* we weight this value by the relative frequencies of the loci with the expected (*P3*⇔*P2*) IG topology (IG loci) and the alternative topology (ALT loci). The *ΔΔΔK* summary statistic is calculated as follows:

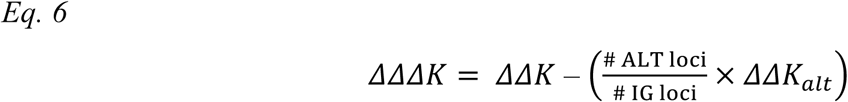

It should be noted that calculation of a *3*×*DIP* correction is only possible when there is at least some mis-assignment/ILS because it relies on the presence of ((*P1, P3*), *P2*) loci. As such, when we applied *3*×*DIP* to genomes simulated with different branch lengths, we were only able to consistently obtain measurements under short-branch conditions (SF < 1.0) where ILS is prevalent (Fig. 6C, F, and I), because these were the only conditions that returned some loci with the relevant topology. Under these short-branch conditions, we found that *3*×*DIP* reduced but did not eliminate the bias observed in *2*×*DIP*. While *ΔΔΔK* was still erroneously positive for the lowest branch length values (Fig. 6F and I), the magnitude of *ΔΔΔK* was less than that of *ΔΔK.*

We further explored bias in *2*×*DIP* and *3*×*DIP* by simulating short branch trees (with SF of 0.1, 0.2, and 0.3) across a range of *p(P3⇒P2*) values. We first applied omniscient *2*×*DIP* to give context to the bias introduced during assignment. As expected, omniscient *2*×*DIP* yielded negative *ΔΔK* values for all replicates in which *p(P3⇒P2*) < 0.5 (Fig. 7A). Consistent with the bias observed in Fig. 6, standard (non-omniscient) *2*×*DIP* yielded erroneously positive *ΔΔK* values, especially for the shortest branch length conditions (Fig. 7B). *3*×*DIP* reduced the bias, only yielding erroneously positive *ΔΔΔK* values for the highest *p(P3⇒P2*) values and the shortest branch length conditions (Fig. 7C). We also tested the performance of *DIP* in a situation in which ILS has occurred but not IG (*pIG*=0; SF=0.1) (Fig. S3). Despite the lack of true IG in these simulations, *1*×*DIP* produced a profile consistent with *P3⇒P2* IG (Fig. S3B), although the relative positions of *ΔK*_*23*_, *ΔK*_*12*_, and *ΔK*_*13*_ distributions differed from the pattern in Fig. 3C. *2*×*DIP* also significantly indicated *P3⇒P2* IG (Fig. S3C), but *3*×*DIP* produced a *ΔΔΔK* that was not significantly different than zero, again indicating that *3*×*DIP* is less prone to falsely indicating *P3⇒P2* IG. Together, these results indicate that *3*×*DIP* is the most robust of the three tests.

**Fig. 7.**
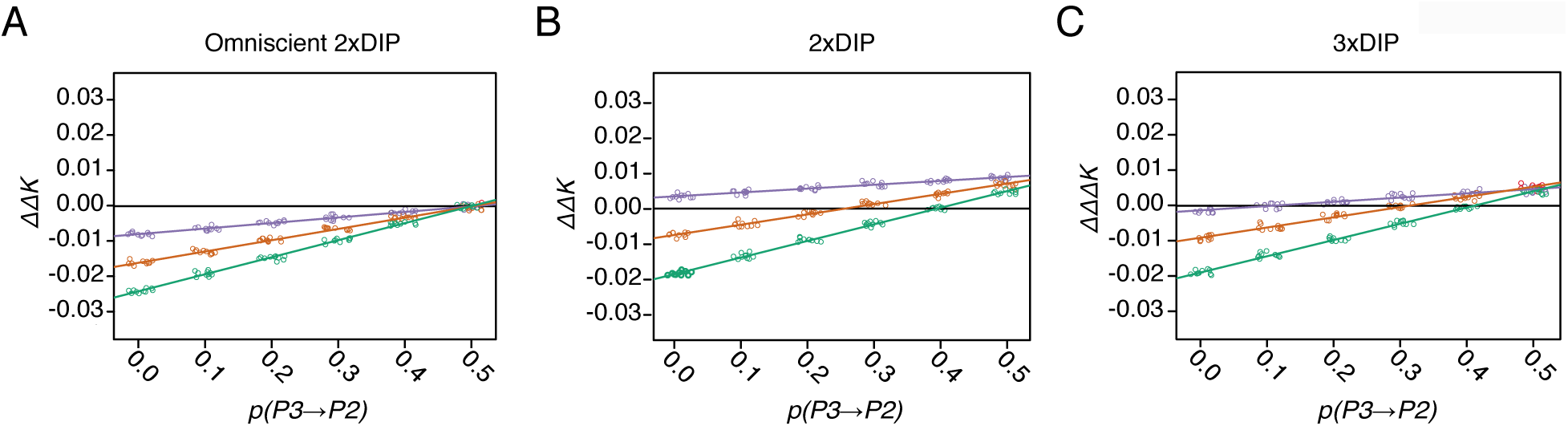
Characterization of *DIP* bias under short branch conditions. Genomes were simulated with different values of *p(P3⇒P2*) (x axis) and different branch length scaling factors (SF) (point colors). See Fig. 6 for description of SF. Purple, SF = 0.1; Orange, SF = 0.2; Green, SF = 0.3. As in Fig. 6, Omniscient *2*×*DIP* (**A**), standard *2*×*DIP* (**B**), and *3*×*DIP* (**C**) were performed. Ten replicate genomes were analyzed for each condition. *pIG* = 0.5 was used for all simulations.

### Analysis of hominin IG

To understand the performance of *DIP* on empirical data, we applied *DIP* to existing genomic data. We focused on IG that occurred between Neanderthal and a modern human European lineage (Green et al. 2010; Prüfer et al. 2014). Using a five-taxon application of the *D-*statistic that made use of the phylogenetic position of multiple modern African populations, a previous study (Green et al. 2010) determined that unidirectional IG occurred Neanderthal⇒European lineages. We applied *DIP* to chromosome one from a Neanderthal sample, a Denisovan sample, two modern human (San [African] and French [European]) samples, and the chimpanzee reference genome. The availability of a Denisovan sample allowed us to infer *DIP* in two different ways using two different taxon sampling schemes (TSS1 and TSS2) (Fig. 8A and F). For both TSSs, there were three gene tree topologies present (Fig. 8B and I), indicating the possibility of mis-assignment due to phylogenetic error and ILS.

**Fig. 8.**
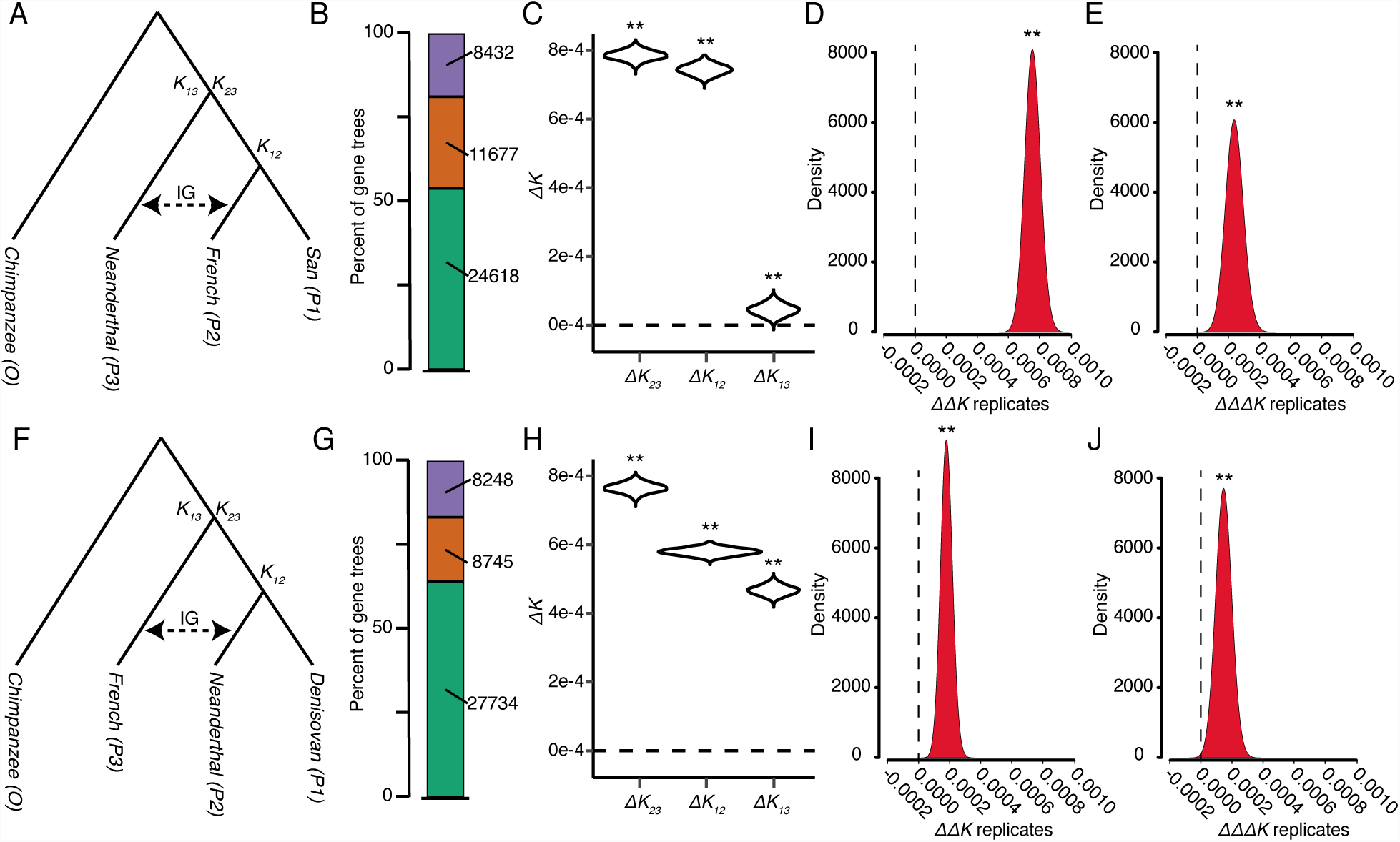
*DIP* analysis of hominin introgression. *DIP* was performed on whole-chromosome alignments of chromosome 1 using two different taxon sampling schemes (TSS). (**A**) Depiction of the samples used in TSS1. (**B**) Neighbor-joining gene-tree topologies from individual loci. (San.,French),Nean.), green; (French, Nean.),San), orange; (San, Nean.),French), purple. (**C-E**) Results from *1*×*DIP* (**C**), *2*×*DIP* (**D**), and *3*×*DIP* (**E**) applied to TSS1 alignment. (**F**) Depiction of the sampled used in TSS2. (**G**) Neighbor-joining gene-tree topologies from individual loci. (Deni.,Nean.),French), green; (Nean.,French),Deni.), orange; (Deni.,French),Nean.), purple. (**H-J**) Results from *1*×*DIP* (**H**), *2*×*DIP* (**I**), and *3*×*DIP* (**J**) applied to TSS2 alignment. ** indicates significant departure from 0 (*p* < 0.01).

Using TSS1, *1*×*DIP* yielded a profile indicating the presence of at least some bidirectional IG (Fig. 8C), a scenario which was not ruled out by (Green et al. 2010). However, it should be noted that, while *ΔK*_*12*_, *ΔK*_*13*_ were both significantly positive, the *ΔK*_*13*_ was much closer to zero, which would indicate a substantial asymmetry towards Neanderthal⇒French IG. *2*×*DIP* and *3*×*DIP* indicated significantly positive *ΔΔK* and *ΔΔΔK,* respectively (Fig. 8D and E), consistent with asymmetric IG in the Neanderthal⇒French direction. However, when we applied *DIP* to TSS2, we saw contradictory results. While, *1*×*DIP* again indicated the presence of bidirectional IG, although without the near-zero *ΔK*_*13*_ (Fig. 8H), *2*×*DIP* and *3*×*DIP* yielded positive *ΔΔK* and *ΔΔΔK,* respectively (Fig. 8I and J). *2*×*DIP* and *3*×*DIP* from TSS2 would indicate French⇒Neanderthal IG. While IG from modern humans has been inferred in other Neanderthal samples (Kuhlwilm et al. 2016), it is at odds with findings from TSS1 and Green et al. (2010).

To understand this discrepancy and put our empirical analyses in the context of our simulations, we plotted distributions of divergence estimates (*K*_*23*_, *K*_*12*_, *K*_*13*_) calculated from two simulated genomes and the TSSs used for the empirical analysis. The empirical distributions display a wider spread than the simulated distributions, potentially introducing noise into the empirical analysis. Importantly, empirical data also show reduced levels of divergence, even compared to the dataset simulated with the shortest branch lengths (SF = 0.1). This suggests that the biasing factors explored above could be even more at-play in the hominin analysis (see Discussion).

## DISCUSSION

### Intended applications of DIP

Our simulation analyses provide a proof-of-principle that divergence data can be used to polarize IG in a four-taxon context, narrowing the methodological gap between our ability to identify IG and our ability to determine the direction of gene transfer. It should be noted that *DIP* is not designed to replace existing methods and act as a frontline test of whether IG has occurred. Instead, we recommend cases of IG first be confidently identified with existing tools (Huson et al. 2005; Than et al. 2008; Green et al. 2010; Durand et al. 2011; Martin et al. 2015; Pease and Hahn 2015; Stenz et al. 2015; Rosenzweig et al. 2016). In these cases, *DIP* can then be used to polarize the direction of IG, a critical step toward interpreting the biological implications of IG. As we have shown above, *DIP* has the potential to distinguish unidirectional and bidirectional IG and, in cases of bidirectionality, to test for asymmetry between the two directions.

While there are population genetic (Schrider et al. 2018) and five-taxon phylogenetic (Green et al. 2010; Pease and Hahn 2015) methods capable of polarizing IG, *DIP* offers the ability to detect asymmetric IG in both directions using a four-taxon context. This will be valuable because very little is known about the extent of reciprocal exchange that occurred during even well-studied IG events (Green et al. 2010; Kuhlwilm et al. 2016), a deficit that likely stems from an absence of sensitive tools. Another group (Hibbins and Hahn, In Review) has recently proposed an approach that overlaps with *DIP*. They introduce a statistic, *D*_2_, which is conceptually similar to *ΔK*_*13*_ described here. As such, non-zero values of *D*_2_ indicate the presence of *P2⇒P3* IG (B*⇒*C by their nomenclature). *DIP* goes further than this approach because it also uses *ΔK*_*12*_ to test for IG in the opposite direction and *ΔΔK* to determine the predominant direction of IG. The primary focus of the recent work by Hibbins and Hahn (In Review), is the development of another statistic, *D*_1_, that assesses the timing of introgression relative to speciation events and can be used in assessing possible cases of homoploid hybrid speciation. This is an elegant application of the same type of divergence-based logic that underlies *DIP* to a biological question that cannot currently be addressed with our method. We suggest that further improvements in polarizing IG can be made by combining the explicit coalescent-based modeling of Hibbins and Hahn with the more comprehensive summary provided by *1*×, *2*×, and *3*×*DIP*.

### Bias in DIP

It should be noted that the simulation branch length parameters used in Fig. 3 and Fig. 5 resulted in gene trees with relatively deep divergences. These branch lengths were chosen because they emphasize differences in divergence and minimize potential biasing factors, thus providing the clearest view of the general properties of *DIP.* However, it has been shown that timing of population divergence is an extremely influential parameter in IG analyses (Durand et al. 2011; Martin et al. 2015; Zheng and Janke 2018). This is true, in part, because the length of internal branches is directly related to the extent of ILS that occurs (Maddison and Knowles 2006). Short branches lead to increased ILS (Degnan and Rosenberg 2013), which can mimic IG and introduce noise into IG analyses. Coalescent simulations, such as those that we performed, capture this phenomenon (Hudson 2002; Degnan and Rosenberg 2009), introducing discordant gene trees at a rate dependent on branch length parameters.

Population divergence is additionally important for *DIP* for a more intuitive reason; the magnitude of the *ΔK* measurements, which are the cornerstone of *DIP,* are directly proportional to the length of internal branches, meaning that *DIP* gains power to differentiate between alternative hypotheses as branches are lengthened. Finally, there is a disparity in the accuracy of topology assignment for loci introgressed *P3⇒P2 vs.* the opposite direction (Zheng and Janke 2018). This disparity stems from the fact that the internal branch on *P2⇒P3* IG gene trees are shorter than the same branch on *P3⇒P2* IG gene trees, making for fewer diagnostic synapomorphies by which to infer the IG topology. This disparity is most pronounced under conditions in which phylogenetically informative synapomorphies are scarce (i.e. short branch lengths). The specific disparity between genes introgressed in each direction is especially problematic for *DIP* because it is likely to introduce a directional bias, favoring inference of *P3⇒P2* IG. All of the above properties lead to challenges at the stage of assigning loci as SP *vs.* IG loci.

For the above reasons, we performed parameter scans to explore the influence of branch length. We found that *2*×*DIP* performs as expected when the assignment step is bypassed in omniscient mode (Fig 6A, D and G) but bias at short branch lengths arises when SP and IG loci must be classified directly based on the data (Fig. 6B, E, and H). Thus, directional bias arises from error at the assignment stage. Of course, when working with empirical datasets, omniscience about origins and the effects of IG *vs.* ILS on individual loci is not possible. As such, assignment error may be unavoidable, so we sought to develop a strategy to correct for bias that arises from mis-assignment, leading to the development of *3*×*DIP.* A benefit of *3*×*DIP* is that it is applicable under the conditions in which bias is most pronounced. Following the logic of the *D-*statistic (Green et al. 2010), *3*×*DIP* is based on the expectation that ILS is equally likely to produce the two topologies that conflict with the species tree: (*P1*(*P2,P3*)) and (*P2*(*P1,P3*)). Therefore, under the assumption that there has been no IG between *P3* and *P1*, the number of “ALT loci”, which are defined by having the (*P2*(*P1,P3*)) topology, provides an estimate for the number of identified “IG loci” that were actually the result of ILS. Accordingly, *3*×*DIP* applies a correction for ILS that is proportional to the frequency of these ALT loci. We found that *3*×*DIP* reduces directional bias at short branch lengths (Fig. 6C, F, and I; Fig. 6) and does not provide false positive results in the complete absence of IG (Fig. S3). These results indicate that *3*×*DIP* is a step toward overcoming directional bias; however, bias persisted for the shortest branch length simulations, meaning that there are biological scenarios in which *3*×*DIP* is not free from bias.

Fully overcoming bias introduced into IG analyses by assignment error represents a future goal for the field. With current implementations of *DIP*, inferences of IG in the *P3⇒P2* direction should be viewed with caution, especially in taxa with very recent divergence times. On the other hand, it can be viewed as a conservative test for *P2⇒P3* IG, so identification of IG in that direction can be interpreted as a much more confident prediction. As suggested above, further progress in this area may come through more complex models that explicitly include ILS (Hibbins and Hahn, In Review).

There are also unexplored factors that should be considered when implementing *DIP* because our simulations were run under simplifying assumptions such as random mating, constant population size, and a single bout of instantaneous IG solely between *P3* and *P2*. Violation of these assumptions in natural populations (Eriksson and Manica 2012; Prüfer et al. 2014; Kuhlwilm et al. 2016; Slon et al. 2018) may introduce additional sources of bias, which should be investigated in future studies with more complex simulation scenarios.

### DIP performance on empirical data

We chose hominin IG as a test case because it is one of the most famous and best-studied examples of IG. An additional benefit is that the sampling in the group is dense; several modern human samples as well as samples from ancient Neanderthal and Denisovan tissues are available. A benefit of this dense taxon sampling is that previous studies have been able to apply five-taxon statistics to polarize IG, leading to the conclusion that “all or almost all of the gene flow detected was from Neandertals into modern humans” (Green et al. 2010). However, more recent analyses of additional archaic samples from different parts of the hominin geographical range also indicated IG in the opposite direction (Kuhlwilm et al. 2016) as well as mating between Neanderthals and Denisovans (Slon et al. 2018).

An additional benefit of dense hominin taxon-sampling is that the phylogenetic placement of samples allows us to analyze the same IG event with four-taxon statistics from two different angles. We devised a TSS in which Neanderthal and a modern human acted as *P3* and *P2*, respectively (TSS1, Fig. 8A) as well as one in which the roles were reversed (TSS2, Fig. 8F). Importantly, these TSSs allowed us to evaluate whether the directional bias described above was strong enough to outweigh the true signature from IG. *DIP* returned contradictory results for TSS1 and TSS2. In both cases, *2*×*DIP* and *3*×*DIP* favored *P3⇒P2* IG, despite the identity of *P3* and *P2* being reversed in the two cases. The fact that both analyses sided with the directional bias we documented above, suggests that bias may be outweighing the IG signature. This is consistent with the observation that hominin divergence is lower than even our shortest simulated branch lengths (Fig. S4), suggesting that biasing factors are strong enough to bias even *3*×*DIP.* It is worth noting, however, that the magnitude of *ΔΔK* and *ΔΔΔK* from TSS1 is higher than that from TSS2, meaning the signal favoring Neanderthal⇒French IG (the expected direction) is stronger than the signal in the opposite direction.

Our general takeaway from analysis of hominin data is that, like all IG analysis tools, there are limits to the conditions under which *DIP* can be reliably applied. Although *3*×*DIP* represents a step in the right direction, in the case of hominin IG, the level of ILS swamps out the signal of IG. We suggest that incorporating an alternative means of assigning introgressed loci, such as *f*_*d*_ (Durand et al. 2011; Martin et al. 2015), may yield more reliable results when ILS is prevalent, representing an area of future work. For the time being, *DIP* will be most reliable in cases of IG that occurred at more ancient time scales (Forsythe et al. In Review; Dasmahapatra et al. 2012; Fontaine et al. 2015).

## METHODS

### Resource availability

URLs for downloading previously published data are provided in place in the following sections. Scripts for reproducing the analyses in this study are available at: https://github.com/EvanForsythe/DIP. Also included are *R* scripts for performing *DIP* on genomic data. All scripts are callable from the command line. Users have the choice of inputting either whole chromosome alignments, which will be divided into single window (i.e. locus) alignments in preparation for *DIP*. Alternatively, *DIP* takes single-locus alignments, bypassing the window partitioning step. *DIP* outputs descriptive statistics and PDF figures similar to Fig. 8.

### Simulations of sequence evolution

We generated whole genome alignments in which IG has occurred in some (but not all) loci, and in which donor and recipient taxa for each introgressed locus are known. To accomplish this, we simulated sequence evolution of loci 5000 nucleotides in length in a four-taxon system (three in-group taxa, *P1, P2*, and *P3* and an outgroup, *O*) (Fig. 1). All simulations were performed with *ms* (Hudson 2002) and *seq-gen* (Rambaut and Grassly 1997) implemented in *R* v3.5.0 with *phyclust* v0.1-22 (Chen 2011) similar to (Martin et al. 2015). A portion of the loci were simulated to have evolved along a path of simple speciation. In the absence of ILS, the gene trees for these loci should match the speciation history, ((*P1,P2*)*P3*)*O*) (Fig. 1A). These loci, denoted as SP loci, were simulated with the following *R* commands:

~~~
ret.msSP<-ms(nsam = 4, nreps = 1, opts = “-T -t 50 -I 4 1 1 1 1 -ej 4 2 1 -ej 8 3 1 -ej 12 4 1 -r 5 5000”)

seqsSP<-seqgen(opts = “-mHKY -l5000 -s 0.01”, newick.tree = ret.msSP[3])
~~~

Loci with instantaneous unidirectional IG occurring between *P2* and *P3* (IG loci) were also simulated. IG trees (transferred in either direction) will have the topology, (*P3,P2*)*P1*)*O*), and thus differ from the species tree. The direction of IG for an individual locus was indicated by ‘donor taxon’ and ‘recipient taxon’ as in the following *R* command:

~~~
ret.msIG <-ms(nsam = 4, nreps = 1, opts= “-T -t 50 -I 4 1 1 1 1 -ej 4 2 1 -ej 8 3 1 -ej 12 4 1 -es 2 <recipient taxon> 0.4 -ej 2 5 <donor taxon> -r 5 5000”)

seqsIG<-seqgen(opts = “-mHKY -l5000 -s 0.01”, newick.tree = ret.msIG[3])
~~~

We replicated the above commands for SP and IG loci to create datasets representing simulated ‘whole-genome alignments’ composed of a total of 5000 loci (Fig. S1). We define the proportion of all loci in the genome resulting from simulated IG in either direction as *pIG* and the proportion of introgressed genes that were transferred in the *P3⇒P2* direction as *p(P3⇒P2).* Since a single locus can only be transferred in one direction or the other, the proportion of loci transferred in the *P2⇒P3* direction, *p(P2⇒P3),* is 1 - *p(P3⇒P2).* Whole genome alignments with known values of *p(IG*) and *p(P3⇒P2*) were used to test the performance of *DIP*. We performed parameter scans by simulating genome alignments with varying combinations of *p(IG*) and *p(P2⇒P3*) (See Fig. S1).

The default branch length parameters used for Fig. 3 and Fig. 5 are *T*_*IG*_=1, *T*_*α*_*=*4, *T*_*β*_=8, and *T*_γ_=12 measured in coalescent units of 4*N* generations (see Fig. 1). To explore the effects of divergence times, we multiplied all branch length parameters by a range of different SF values. For example, SF=0.1 results in the following node depths: *T*_*IG*_=0.1, *T*_*α*_*=*0.4, *T*_*β*_=0.8, and *T*_*γ*_=1.2. For parameter scans involving branch lengths, we generated point estimates of *ΔΔK* and *ΔΔΔK* from ten replicate genomes for each condition.

### Assignment of SP and IG loci

The first step in all versions of *DIP* is sorting loci to isolate the loci that were introgressed and those that follow the species branching order (i.e. topology assignment). Using simulated data affords us omniscience at this step (i.e. we know whether each locus was originally simulated as introgressed or not). However, unless specifically stated, we did not make use of the known history of simulated loci. Instead, *DIP* infers the IG status of loci based on the topology of a neighbor joining gene tree inferred for each locus using *Ape* v5.2 (Paradis et al. 2004). Loci displaying the ((*P1,P2*)*P3*)*O*) topology are marked as speciation loci (SP loci). Loci displaying the ((*P2,P3*)*P1*)*O*) topology are designated as introgressed loci (IG loci). Any loci displaying the alternative topology, ((*P1,P3*)*P2*)*O*), which are not produced by speciation or IG, are omitted from *1*×*DIP* and *2*×*DIP* but used by *3*×*DIP* to calculate a correction factor (see below).

### Inferring IG directionality with 1×*DIP*

We calculated the pairwise divergences, *K*_*23*_, *K*_*12*_, and *K*_13_ (as indicated in Fig. 1A) for each IG and SP locus using the *dist.dna* command from the *Ape* package with default settings. Pairwise divergences, *K*_*23*_, *K*_*12*_, and *K*_*13*_ are named for the taxa involved in the distance calculation. For example, *K*_*23*_ measures the divergence of *P2* and *P3* (see Fig. 1). *ΔK*_*23*_, *ΔK*_*12*_, and *ΔK*_*13*_ were calculated based on difference in mean *K* values between SP and IG loci as shown in *Eqs. 1-3*. To test for significance, bootstrapped distributions were obtained by resampling (with replacement) loci from the genome to achieve genome alignments equal in number of loci to the original genome alignment. 1000 such replicates were performed, recalculating *ΔK*_*23*_, *ΔK*_*12*_, and *ΔK*_*13*_ for each replicate. *P*-values for the significance of *ΔK* values were calculated as the proportion of replicates for which *ΔK* ≤ 0.

### Inferring IG directionality with 2×DIP and 3×DIP

*ΔΔK* was calculated from *ΔK*_*12*_, and *ΔK*_*13*_ described in *Eq. 3*. The bootstrap resampling scheme described in the previous paragraph was used to assess the significance of *2*×*DIP. ΔΔK* was calculated for each replicate and *p-*values were obtained from the proportion of replicates for which *ΔΔK* overlapped zero (multiplied by two for a two-sided test). Like *2*×*DIP, 3*×*DIP* makes use of *ΔΔK* to indicate the directionality of IG. However, *3*×*DIP* also introduces *ΔΔK*_*alt*_, *w*hich is calculated according to *Eq. 5. ΔΔΔK* is obtained from the difference between *ΔΔK* and *ΔΔK*_*alt*_ (see *Eq. 6*). As for *ΔΔK* above, significance of *ΔΔΔK* is obtained from resampled whole genomes alignments.

### Hominin data analysis

To generate whole-chromosome alignments from the hominin dataset for *DIP*, genome resequencing data for two Neanderthal, one Denisovan, and two modern human samples from (Prüfer et al. 2014) were downloaded from http://cdna.eva.mpg.de/neandertal/. VCF files were downloaded for chromosome 1 for each species. The human reference genome (hg19) (International Human Genome Sequencing Consortium 2001), which was originally used for read mapping during the creation of VCF files, was obtained from http://hgdownload.cse.ucsc.edu/goldenPath/hg19/. The following procedures were performed for each sample.

Structural variation (indel) information was trimmed from VCF files, using *VCFtools* (Danecek et al. 2011) and *Tabix* (Li et al. 2009) with the following commands:

~~~
vcftools --gzvcf Chrom1_with_indels.vcf.gz --remove-indels --recode --recode-INFO-all --out Chrom1_SNPs_only.vcf

bgzip Chrom1_SNPs_only.vcf

tabix -p vcf Chrom1_SNPs_only.vcf.gz
~~~

Whole-chromosome consensus sequence was extracted from VCF files using *BCFtools* (Li et al. 2009) with the command below. For heterozygous sites, by default *bcftools consensus* applies the alternative variant (i.e. the variant that does not match the reference genome) to the consensus sequence for the given sample (see https://samtools.github.io/bcftools/bcftools.html).

~~~
cat hg19_chrom1.fa | bcftools consensus Chrom1_SNPs_only.vcf.gz > Chrom_1_consensus.fa
~~~

We used the reference chimpanzee genome (PanTro5) (The Chimpanzee Sequencing Consortium 2005) as an outgroup. We downloaded a MAF alignment of chromosome one from PanTro5 and hg19 from: http://hgdownload.cse.ucsc.edu/goldenpath/hg19/vsPanTro5/axtNet/. We converted this file to a FASTA file using Galaxy tools (Afgan et al. 2018) available at https://usegalaxy.org/. Finally, the consensus sequence from each hominin samples and chimpanzee was concatenated into a whole-chromosome multiple sequence alignment in FASTA format. This five-taxon alignment was pruned to contain four taxa according to each TSS (see Fig. 8) and then used as input to *DIP.*

## ACKNOWLEDGMENTS

This work was funded by NSF grant MCB-1733227 to D.B.S. as well as NSF grant IOS-1444490 to M.A.B. We thank M.J. Sanderson, R.A. Mosher, A.D.L Nelson, K. Dew-Budd, K. Palos, A.E. Baniaga, and S.M. Lambert for helpful discussion.

## SUPPLEMENTAL INFORMATION

**Fig. S1.**
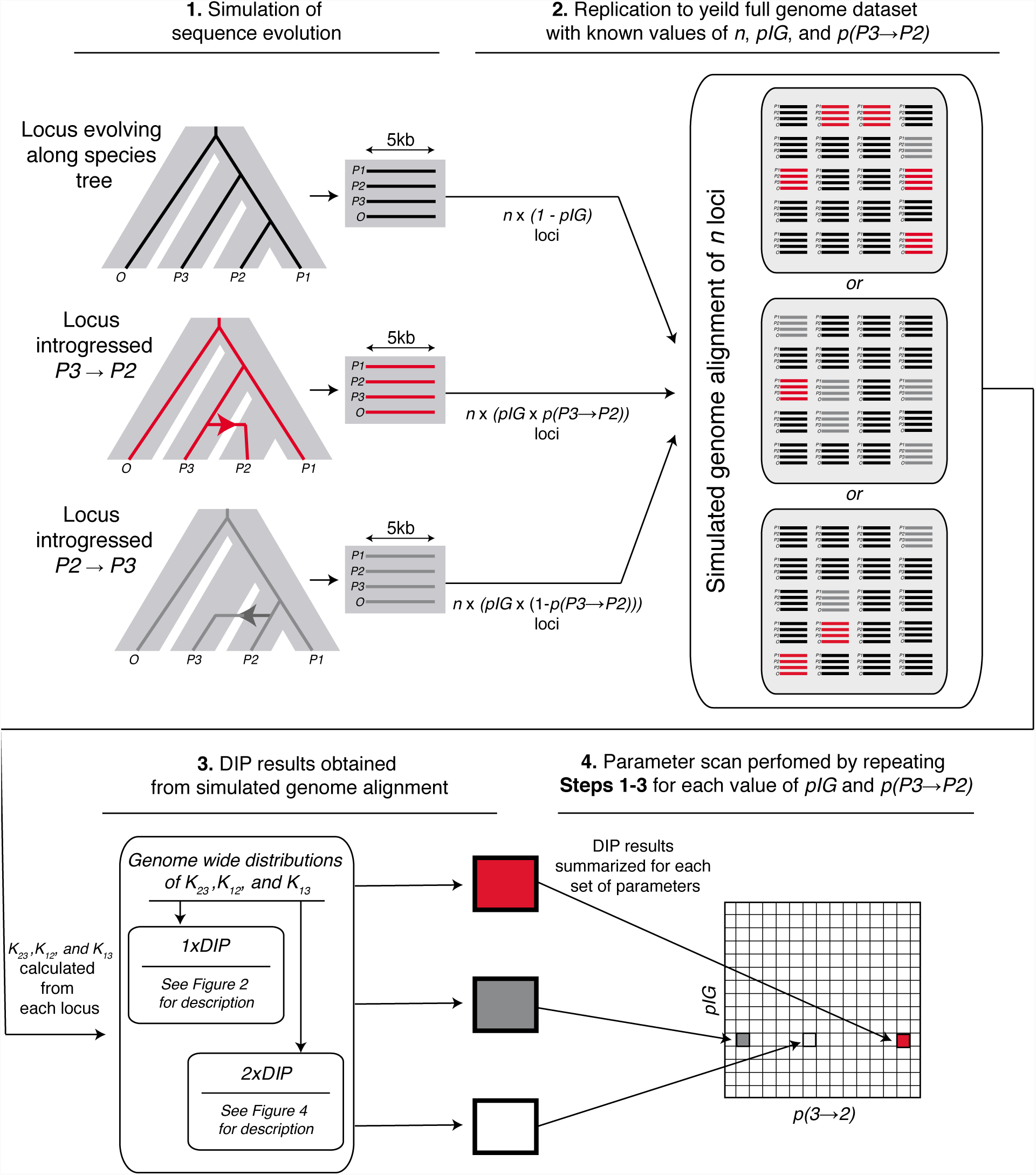
Schematic of the workflow used to simulate IG across a genome and perform *DIP*. **(1)** Each locus is evolved along the species tree or along a path of IG and used to generate a 5kb alignment using *ms* and *seq-gen* similar to (Martin et al. 2015). **(2)** Step 1 was repeated to yield a full genome of *n=*5000 loci in which *n* x *p(IG*) loci were introgressed and the remaining loci evolved along the species tree. For example, a genome in which half of all genes were not transferred while the other half were transferred *P3⇒P2* would be generated with: *n=5000, pIG* = 0.5, *p(P3⇒P2*) = 1.0. **(3)** Different steps in the *DIP* pipeline are performed on the simulated genome. **(4)** Steps 1-3 are repeated for each combination of *pIG* and *p(P3*⇒*P2*). Each pixel in a parameter scan graph represents one or more runs of Steps 1-3.

**Fig. S2.**
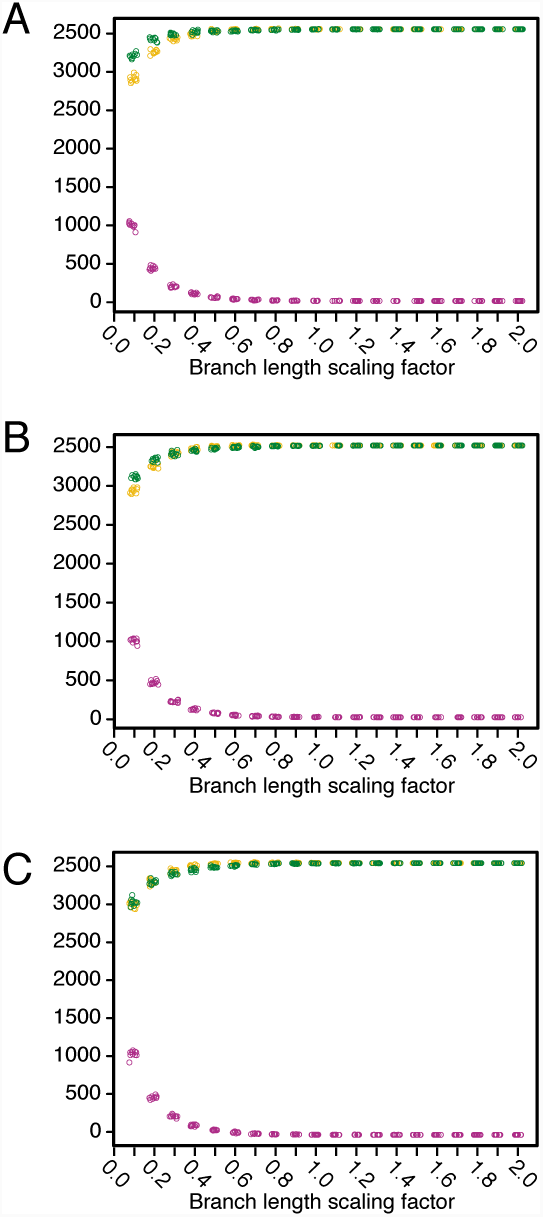
Gene tree topologies inferred from simulated genomes. Gene tree counts for genomes simulated with different branch lengths (x-axes) and *p(P3⇒P2*) values of 0.6 (**A**), 0.5 (**B**), and 0.4 (**C**). Each point represents the number of trees displaying a given topology from a replicate genome. ((*P1,P2*),*P3*), *orange;* ((*P2,P3*),*P1*), *green;* ((*P1,P3*),*P2*), *purple.* These same simulated genomes were analyzed in Fig. 6.

**Fig. S3.**
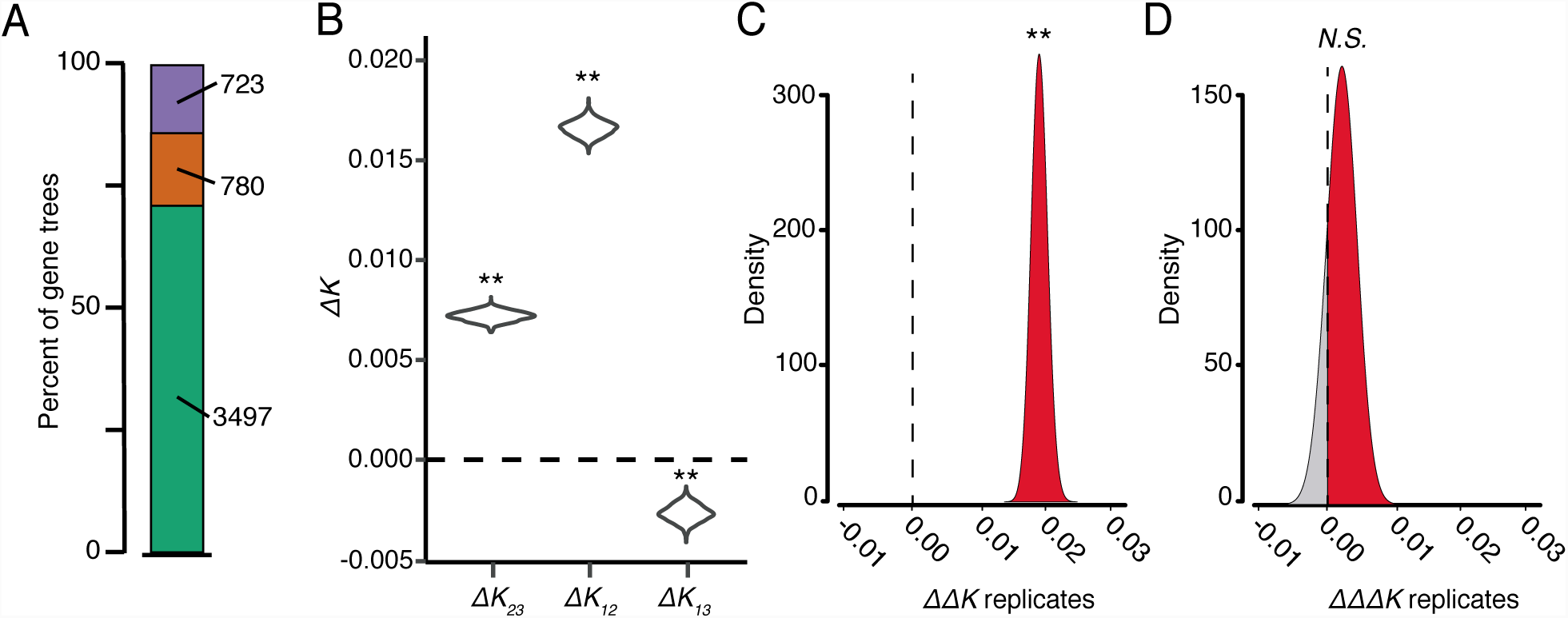
*DIP* analysis of a genome with incomplete lineage sorting but no introgression. A genome alignment was simulated with *pIG* set to zero using the scaling factor 0.1 (see Fig. 1 and Fig. 6). Therefore, all loci with topologies that conflict with species tree are the result of ILS and not IG (**A**) The topologies of neighbor joining trees inferred from 5000 simulated loci. ((*P1,P2*),*P3*), *green;* ((*P2,P3*),*P1*), *orange;* ((*P1,P3*),*P2*), *purple.* (**B-D**) 1×*DIP* (**B**), 2×*DIP* (**C**) and 3×*DIP* (**D**) analysis of the genome alignment.

**Fig. S4.**
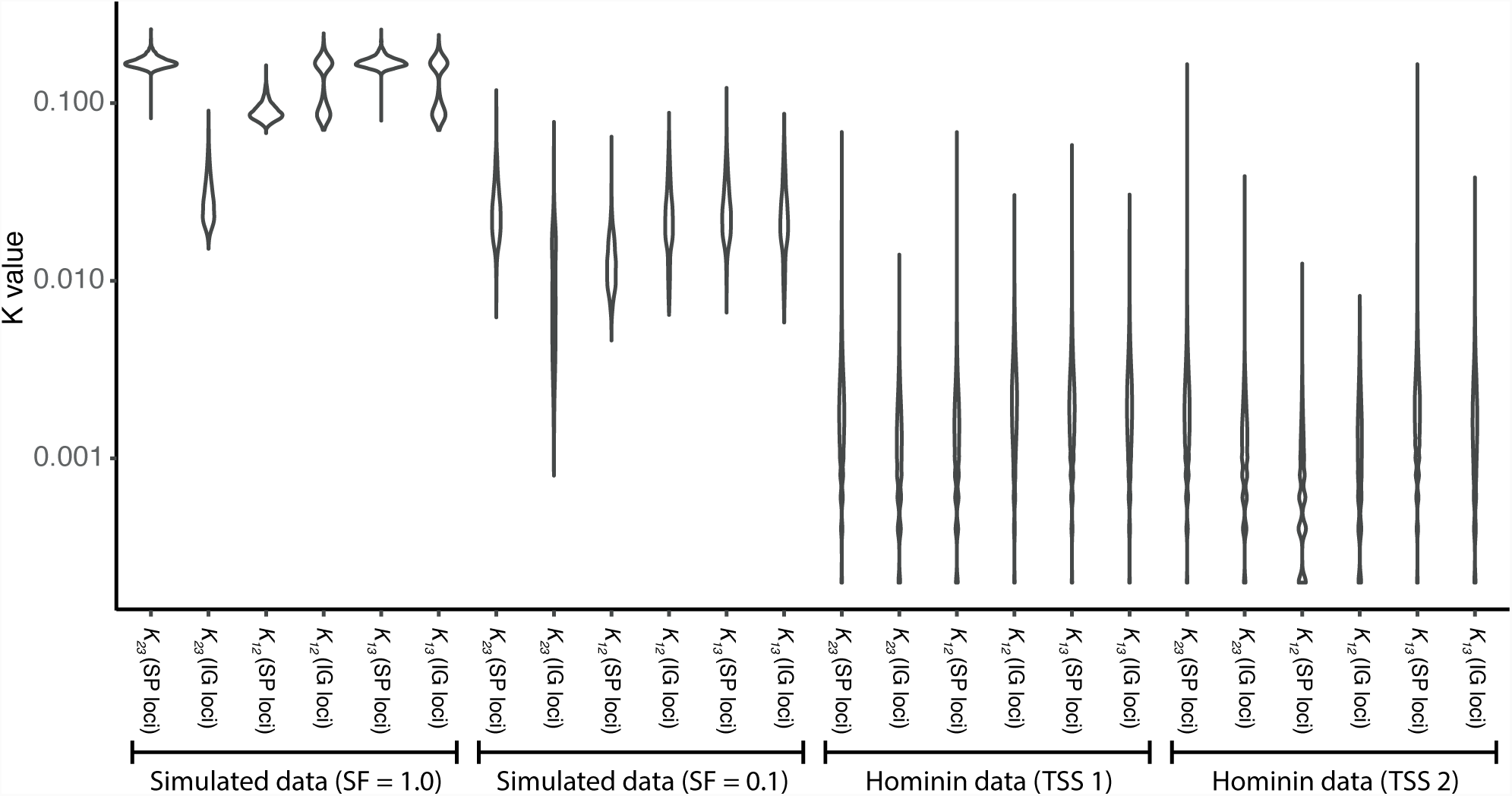
Sequence divergence measures from simulated and Hominin data. Violin plot showing distributions of pairwise divergence values for inferred SP and IG loci (see Fig. 1 and 2). Both simulated datasets were simulated with *pIG*=0.5 and *p(P3⇒P2*)=0.5.

